# Translation-dependent and independent mRNA decay occur through mutually exclusive pathways that are defined by ribosome density during T Cell activation

**DOI:** 10.1101/2020.10.16.341222

**Authors:** Blandine C. Mercier, Emmanuel Labaronne, David Cluet, Alicia Bicknell, Antoine Corbin, Laura Guiguettaz, Fabien Aube, Laurent Modolo, Didier Auboeuf, Melissa J. Moore, Emiliano P. Ricci

## Abstract

mRNA translation and degradation are strongly interconnected processes that participate in the fine tuning of gene expression. Particularly, targeting mRNAs to translation-dependent degradation (TDD) could attenuate protein expression by making any increase in mRNA translation self-limiting. However, the extent to which TDD is a general mechanism for limiting protein expression is currently unknown. Here we describe a comprehensive analysis of basal and signal-induced TDD in mouse primary CD4 T cells. Our data indicate that most cellular transcripts are decayed to some extent in a translation-dependent manner, both in resting and activated cells. Our analysis further identifies the length of untranslated regions, the density of ribosomes and the GC content of the coding region as major determinants of TDD magnitude. Consistent with this, all transcripts that undergo changes in ribosome density upon T cell activation display a corresponding change in their TDD level. Surprisingly, the amplitude of translation-independent mRNA decay (TID) appears as a mirror image of TDD. Moreover, TID also responds to changes in ribosome density upon T cell activation but in the opposite direction from the one observed for TDD. Our data demonstrate a strong interconnection between mRNA translation and decay in mammalian cells. Furthermore, they indicate that ribosome density is a major determinant of the pathway by which transcripts are degraded within cells.

## Introduction

mRNA degradation contributes to defining steady-state transcript levels, removing aberrant mRNAs through surveillance pathways and also to the dynamic regulation of mRNA abundance in response to cellular cues. mRNA degradation involves a wide variety of effectors and regulatory factors, which for the most part correspond to RNA-binding proteins that are able to recognize specific sequences or structural motifs in the target mRNAs. Similar to most other steps in the gene expression pathway, mRNA degradation is often coordinated with upstream and downstream steps. For example, mRNA translation and decay are strongly interconnected, particularly in the context of mRNA surveillance pathways^1^. Although Translation-Dependent Decay (TDD) pathways were originally identified and studied as the means by which cells rid themselves of aberrant mRNAs (e.g., those containing premature termination codons in the case of non sense mediated decay (NMD), and truncated or prematurely polyadenylated mRNAs in the case of non-stop decay (NSD)), it is now widely recognized that NMD is also a key post-transcriptional regulatory mechanism for physiologically functional mRNAs^2^. Furthermore, other translation-dependent mRNA decay pathways have been discovered that regulate stability of physiologically functional transcripts bearing binding sites for specific RNA-binding proteins (such as Staufen^3^), through specific codon usage^4–13^, or as a consequence of ribosome collisions^14–17^. Co-translational mRNA degradation^18–20^ is therefore emerging as a major decay pathway for functional mRNAs in eukaryotic cells. However the extent to which TDD is a general mechanism for limiting the number of protein molecules made per mRNA molecule, both basally and in response to signaling, is currently unknown. Furthermore, the relationship between TDD and translation-independent mRNA decay (TID) pathways has not yet been evaluated in a global manner.

Transcriptional and post-transcriptional controls are crucial for regulating gene expression, both basally and in response to extracellular cues. One way that protein expression might be attenuated following translational activation is by targeting mRNAs to TDD, thus making any increase in protein expression both transient and self-limiting. When coupled to transcriptional control, TDD might further participate in temporal regulation of gene expression by allowing rapid clearance of transcriptionally repressed genes.

Here we describe a comprehensive analysis of basal and signal-dependent gene expression in primary resting and activated mouse CD4+ T lymphocytes. We performed RNA-Seq, poly(A)-site sequencing and ribosome profiling to monitor RNA levels, 3’-untranslated region (UTR) length and translational efficiency, at the transcriptome-wide level, both before and after activation. We used transcription inhibitors to assess RNA degradation and translation inhibitors to evaluate the prevalence of TDD. This strategy allowed us, for the first time, to discriminate between translation-dependent (TDD) and translation-independent (TID) mRNA degradation in a transcriptome-wide manner. Surprisingly, our data indicate that most unstable mRNAs are decayed at least partially in a translation-dependent manner in both resting and activated T cells. Furthermore, the extent of translation-dependent and independent mRNA degradation is governed by similar *cis*- and *trans*-acting features, albeit in an opposing manner, thus suggesting that both pathways are competing against each other.

## Results

### Activation of primary mouse CD4+ T cells induces profound transcriptome and translatome remodeling

Highly pure primary mouse CD4 T cells (>90% CD3+ CD4+; figure 1a) were obtained by negative selection from mouse spleens and lymph nodes (Figure 1b). These purified cells were then cultured *in vitro* in the absence (Resting) or presence (Activated) of anti-CD3/anti-CD28 antibody-coated beads, mimicking antigen-presenting cells (Figure 1b). Activation was confirmed by strong cell surface expression of the activation markers CD69 and CD25 after 24, 48 and 72 hours, as well as cellular proliferation using CFSE staining (Figure 1c).

**Figure 1.**
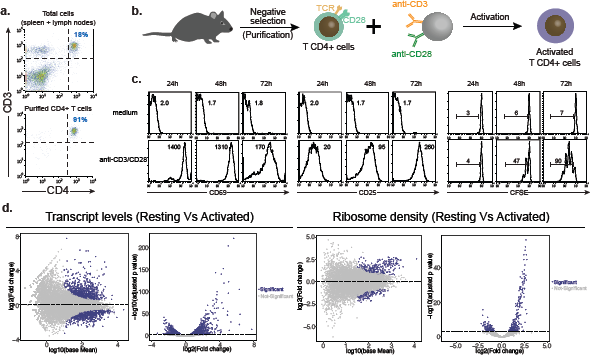
Primary mouse CD4+ T cell purification and activation. **a.** Flow cytometry analysis of CD3 and CD4 surface expression in cells obtained from spleen and lymph nodes before (top panel) and after (bottom panel) purification by negative selection. **b.** Schematic representation of the procedure for purification and activation of primary CD4+ T lymphocytes obtained from C57BL/6J female mice. **c.** Flow cytometry analysis of the cell activation markers (CD69 and CD25) as well as cell division (CFSE) in resting CD4+ T lymphocytes (top panels) or upon activation for 24, 48 and 72 hours in presence of anti-CDS/CD28 magnetic beads. **d.** Differential gene expression analysis for transcript abundance and ribosome density in resting and activated cells.

To enable transcriptome-wide analyses, we prepared whole cell RNA-Seq and ribosome footprinting libraries from three independent biological replicates of T cells after 0 (Resting) and 3 hours of activation (Activated). To obtain gene-specific expression values, we aligned RNA-Seq reads to the GENCODE APPRIS mouse principal transcript reference set^1^. In agreement with the flow cytometry results, anti-CD3/CD28 stimulation significantly altered the T cell transcriptome (Figure 1d and Table S1). In all, 822 transcripts showed a significant increase in expression upon activation and 694 transcripts exhibited decreased expression (Figure 1d, left panel). Gene-ontology analysis indicates that differentially up-regulated transcripts at 3h post activation include pathways related to gene expression regulation such as ribosome biogenesis, translation initiation, tRNA processing, mRNA processing and regulation of RNA polII transcription, as well as pathways related to immune system and response to cytokine (Supplementary Figure 1a and Table S1). Significantly down-regulated transcripts are enriched for ribosomal proteins, protein ubiquitination factors and factors involved in cell cycle, T cell differentiation and RNA splicing among others (Supplementary Figure 1b and Table S1).

Similarly, anti-CD3/CD28 stimulation, altered the translatome of T cells (Figure 1d, right panel). Ribosome profiling revealed hundreds of transcripts displaying significant changes in ribosome density upon T cell activation (Figure 1d, right panel). Those displaying a significant increase in ribosome density (256 transcripts in total) are enriched for ribosomal proteins (78 out of the 80 core ribosomal proteins), ribosome biogenesis factors, RNA splicing factors and factors involved in the cellular response to interleukin-4 (Supplementary Figure 1c and Table S1). Conversely, transcripts displaying a significant decrease in ribosome density (117 transcripts in total) mainly code for kinases, signal transduction factors and transcription factors (Supplementary Figure 1d and Table S1). Thus, activation of T cells results in a profound remodeling of their transcriptome and reveals significant changes in ribosome density across hundreds of transcripts. For some cellular functions, such as ribosome biogenesis and mRNA splicing, changes in transcript levels and ribosome density are coordinated to potentially maximize protein output during activation. By contrast, expression of ribosomal proteins appears to be regulated in a manner where translation up-regulation could buffer some of the observed decrease in transcript abundance upon T cell activation.

### Monitoring mRNA decay in a transcriptome-wide manner

To monitor mRNA decay in resting and activated T cells, we utilized a transcription inhibition strategy. For this, we tested the efficiency of two different transcription inhibitors in actively transcribing activated T cells, 5,6-Dichloro-1-β-D-ribofuranosylbenzimidazole (DRB), which inhibits RNA polymerase II elongation^2^, and Triptolide which irreversibly blocks transcription from RNA polymerase II through targeting of the general transcription factor TFIIH^3^. Because transcription inhibitors induce a global decrease in mRNA abundance within cells, external RNA controls consortium (ERCC) spike-in pool was added to cells during RNA extraction in order to obtain absolute quantification of transcript abundance at each time point upon transcription inhibition. As expected, the relative amount of reads mapping to spike-in RNAs increased upon blocking transcription when compared to those mapping to endogenous mRNAs (Supplementary Figure 2a). However, spike-in normalization of read counts introduced a significant source of technical variability. To avoid this issue, we decided to use spike-in read counts to identify stable endogenous transcripts that exhibited almost no decay upon transcription inhibition over the three-hour time course of our experiments (see methods section), both in resting and activated cells (Supplementary figure 2b). We then used these stable transcripts to normalize abundance of all transcripts from our libraries using the DESeq2 package^4^. Analysis of our datasets allowed us to monitor the fold change in RNA abundance after one and three-hour transcription blockade and to obtain a proxy of decay rates in resting and activated T cells. From these data, triptolide appeared to inhibit transcription more efficiently than DRB at both time points (Figure 2c, left lanes) and was therefore selected to inhibit transcription in most experiments.

**Figure 2.**
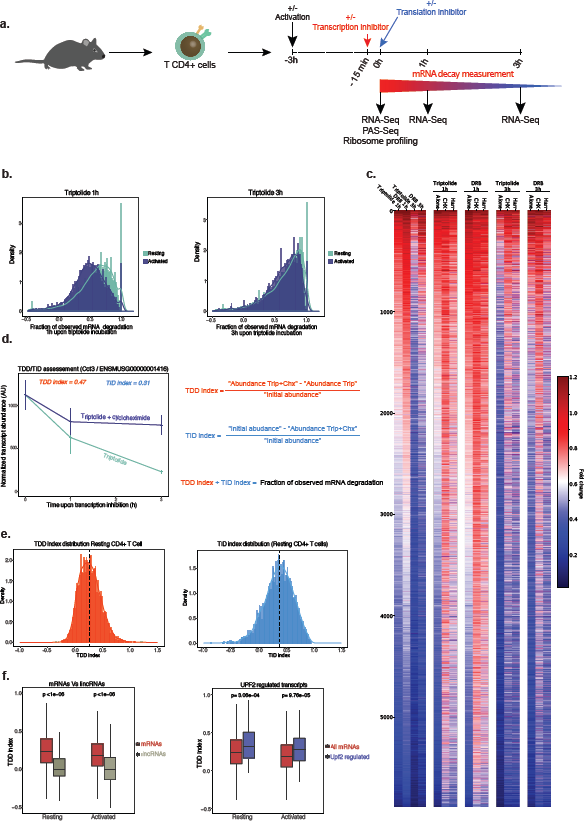
Monitoring translation-dependent and independent mRNA decay in CD4+ T lymphocytes. **a**. Schematic representation of the procedure to monitor overall and translation-dependent mRNA degradation. Purified CD4+ T lymphocytes are activated or not for 3 hours. After this, cells are incubated or not with transcription inhibitors (Triptolide or DRB). 15 minutes following addition of transcription inhibitors, translation inhibitors (cycloheximide or harringtonine) are added to cells. Cells are collected at 0, 1 and 3 hours following transcription inhibition to monitor transcript expression by RNA-sequencing. Ribosome profiling and poly(A)-site sequence (PAS-seq) were also performed at the time 0 h in absence of transcription inhibitors. **b.** Fraction of mRNA degradation in resting and activated T cells 1 and 3 hours after transcription inhibition with triptolide. **c.** Fold change in transcript abundance (compared to the 0 h time point) in activated CD4+ lymphocytes upon blocking transcription only or when blocking both transcription and translation. **d.** Details of the calculation of the TDD and TID indexes illustrated by the expression dynamics of the transcript coding for CCT3 upon incubation with transcription and translation inhibitors. **e.** Distribution of the TDDindex and TIDindex calculated from resting T cells incubated for 3 hours in the presence of triptolide and cycloheximide. **f.** Comparison of the TDDindex (triptolide and cycloheximide) for protein coding transcripts (mRNAs) and long intergenic non-coding RNAs (lincRNAs) (left panel) or for all protein coding transcripts (all mRNAs) and Upf2-regulated transcripts (right panel) in resting and activated T cells. **g.** Distribution of TDDindex against TIDindex values (calculated using triptolide and cycloheximide) in resting T cells.

Comparison of the fraction of observed mRNA degradation (*Observed mRNA degradation*=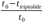) at 1h and 3h upon triptolide addition in both resting and activated T cells (Figure 2b) indicates that mRNA decay is generally more efficient in resting cells as compared to activated cells. This result is consistent with a recent report describing a global stabilization of mRNAs upon CD4+ T cell activation in mice^5^.

### Inhibition of translation stabilizes numerous transcripts in T cells

To assess the extent to which mRNA decay is dependent on translation, we combined transcription and translation inhibitors to obtain mRNA degradation rates in the presence and absence of translation (Figure 2a). To inhibit mRNA translation, we used cycloheximide which blocks elongating ribosomes on the coding sequence, or Harringtonine which blocks the late steps of translation initiation resulting in the run-off of elongating ribosomes and the accumulation of initiating 80S ribosomes at translation start sites. Translation inhibition in primary T cells was validated through metabolic labeling of newly synthesized proteins with [^35^S] methionine in the presence or absence of translation inhibitors (Supplementary Figure 2d). To control for specific biases introduced by each of the transcription inhibitors (DRB or triptolide) and translation inhibitors (cycloheximide or harringtonine), we used them in all possible combinations in activated T cells. As the two transcription inhibitors yielded similar results (Figure 2c), only triptolide was used with either cycloheximide or harringtonine in resting T cells. Briefly, resting or activated T cells were incubated with the above mentioned combinations of inhibitors for different time points (0, 1h or 3h after transcription or transcription plus translation inhibition) and RNA levels were assessed by RNA-Seq, from three independent biological replicates for all conditions. Unexpectedly, results obtained using either DRB or triptolide in activated cells showed that translation inhibition, whether through cycloheximide or harringtonine, led to a global stabilization of mRNAs at both 1h and 3h time points (Figure 2c). These results strongly suggest that translation plays a significant role in mediating degradation of most mRNAs expressed in primary T cells.

To quantify the extent to which overall mRNA decay is dependent on ongoing translation, we calculated a translation-dependent RNA decay index for each time point (TDDindex, Figure 2d). This index corresponds to the difference in absolute transcript abundance between blockade of both transcription and translation and blockade of transcription only, normalized by the initial transcript abundance prior to transcription inhibition (Figure 2d, see TDD index formula). Similarly, we calculated a translation-independent RNA decay index (TIDindex) which measures the difference between transcript abundance at the initial time and after blocking of transcription and translation, normalized by the initial transcript abundance (Figure 2d, see TID index formula). For any given transcript, the sum of the TDDindex and TIDindex corresponds to the fraction of observed mRNA degradation at a given time after blocking transcription. Transcripts whose decay is largely translation-independent have TDDindex values close to 0 and TID values >0, whereas transcripts whose decay is mainly translation-dependent have TDDindex>0 and TIDindex close to 0. Furthermore, TDDindex and TIDindex should be partially independent from each other, the only constraint being that the sum of the TID and TDD index cannot be greater than 1.

TDDindex obtained from the different combinations of transcription and translation inhibitors yielded very similar values with Pearson correlation coefficients ranging from 0.67 to 0.82 (Supplementary figure 2e). This indicates a robust measure of translation-dependent mRNA decay independently of the transcription or translation inhibitor that was used. Similar results were obtained for TIDindex (data not shown). Therefore, for most subsequent analyses, we used the TDDindex and TIDindex obtained from cells treated with triptolide and cycloheximide and, when needed, also showed corresponding results obtained with triptolide and harringtonine as supplementary data.

Distribution of the TDDindex calculated at 3h upon blocking transcription (Figure 2e, left panel) shows a median value of 0.27 (i.e half of the transcripts being degraded at least by 27% at 3h through a translation-dependent mechanism) thus confirming our previous observation that a large fraction of transcripts are degraded, at least partially, in a translation-dependent manner. Only a minor fraction of transcripts displayed high TDDindex values (close to 1) corresponding to highly unstable mRNAs relying almost entirely on translation for their decay (Figure 2e, left panel). In contrast, a larger fraction of transcripts displayed TDDindex values close to 0 or slightly negative, indicating that their degradation is not dependent on translation (Figure 2e, left panel). As a quality control of these observations, we examined the extent of translation-dependent RNA decay of long intergenic non-coding RNA (lincRNA). Because lincRNAs are transcribed by RNA polymerase II, they share many features with protein-coding mRNAs including a 5’ cap structure, excision of introns by the spliceosome and a 3’ poly(A) tail^6^. However, they are mostly not subjected to translation and therefore should not be decayed in a translation-dependent manner. Consistent with this, the TDDindex of lincRNAs (n=585 in resting and n=595 in activated cells) was centered around 0 (Figure 2f, left panel). Conversely, the median TDDindex of transcripts previously identified in mouse thymocytes as regulated by UPF2 through NMD^7^ (n=111 in resting and n=110 in activated cells) was significantly higher than the median TDDindex of all protein-coding mRNAs (Figure 2f, right panel). Our calculated TDDindex thus behave as expected for transcripts whose decay should be either independent (lincRNAs) or dependent (NMD targets) on translation.

Distribution of the TIDindex is overall broader than that of the TDDindex (Figure 2e, right panel) and displays a larger median value of 0.43 (i.e half of the transcripts being degraded at least by 43% at 3h through a translation-independent mechanism). TID is therefore globally preponderant over TDD in defining the extent of mRNA decay in T cells. Interestingly, the global mRNA stabilization observed upon T cell activation (Figure 2b) corresponds mainly to a decrease in the efficiency of TDD rather than TID (Supplementary Figure 2c). Taken together, our results suggest that TDD is involved in the degradation of a large fraction of cellular mRNAs species, although to a less extent than TID, which is the main pathway of mRNA degradation in T cells.

### Functional categories associated with strong and weak TDD and TID transcripts

To identify functional gene ontology categories associated with low or high TDDindex and TIDindex values, we collected GO annotations for all expressed transcripts. Then, for each GO term with at least 10 genes expressed in our samples, we calculated the mean TDDindex and TIDindex and compared them against the mean TDDindex and TIDindex values of the entire transcript population (Figure 3a and supplementary figure 3a). To test for statistical significance between the TDDindex of any given GO term and that of the entire population, we performed permutation tests (see material and methods). This analysis revealed that transcripts with a high TDDindex (i.e. targeted more by TDD) tend to be enriched in core gene expression functions such as ribosome biogenesis, mRNA splicing, tRNA processing, RNA helicase activity and the proteasome complex (Figure 3b). Additionally, categories such as DNA repair and mitochondrial inner membrane were also enriched in transcripts with high TDDindex. In contrast, transcripts with low TDDindex (i.e. targeted less by TDD) appear to be enriched in RNA pol II transcription factors, proteins associated to the plasma membrane, signal transduction, protein kinase activity and ribosomal proteins. Interestingly, while resting T cells share most of their enriched GO terms with that of activated T cells, there are numerous GO categories that are specific to activated T cells, both among high TDDindex and low TDDindex values. This suggests that TDD could play a dynamic role in regulating specific transcript categories upon T cell activation.Regarding translation-independent mRNA decay, some functional categories enriched in low and high TID transcripts showed a mirror image of those observed for translation-dependent mRNA decay. This suggests that some functional gene categories are mostly regulated by one of the two pathways (supplementary figure 3b). For example, transcripts coding for RNA pol II transcription factors and proteins related to stress granules are enriched among high TIDindex and low TDDindex groups. In contrast, transcripts coding for DNA repair and mitochondrial membrane proteins are enriched among low TIDindex and high TDDindex groups.

**Figure 3.**
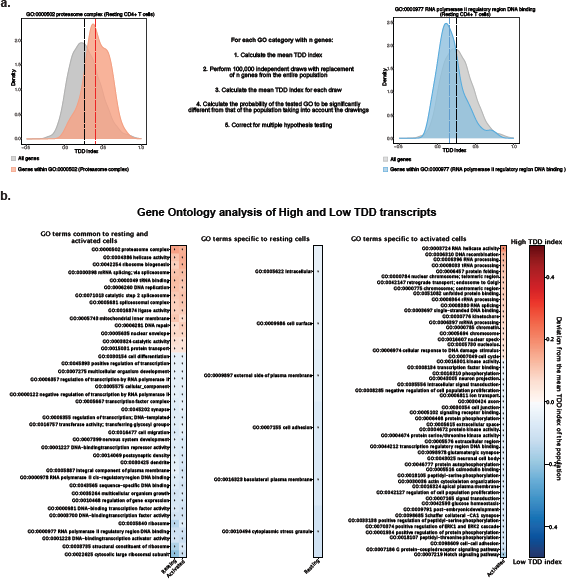
Gene ontology analysis of transcripts displaying high or low TDDindex values. **a. (Left panel)** TDDindex distribution for all expressed genes (grey) or genes from the “Proteasome complex” GO category (red). **(Middle panel)** Scheme describing the procedure to identify GO categories enriched among high and low TDD transcripts. **(Right panel)** TDDindex distribution for all expressed genes (grey) or genes from the “RNA polymerase II regulatory region DNA binding” GO category (blue). **b.** List of GO terms with a median TDDindex that is significantly higher (red) or lower (blue) from the median TDDindex of the entire transcript population that are common to resting and activated T cells (left panel) or specific to resting (middle panel) or activated (right panel) T cells.

Interestingly, transcripts encoding RNA splicing factors appear strongly regulated both by TDD and TID. Conversely, other categories such as mRNAs coding for ribosomal proteins, synapse or protein serine/threonine kinases are enriched in both low TDD and TID transcript groups, therefore indicating that they correspond to stable transcripts. Finally, GO categories related to the immune response (such as innate immune response, cellular response to lipopolysaccharide and immune system process) are mostly regulated through a translation-independent mRNA decay pathway.

Taken together, our data show that different classes of functional categories rely either on TDD or TID as their main degradation pathway while some rely on both. Interestingly, whereas transcripts coding for ribosomal proteins appear as stable, those coding for ribosome biogenesis factors rely mainly on TDD for their decay. This suggests a potential feedback loop where ribosome biogenesis factor availability could be controlled in a translation-dependent manner.

### The length of untranslated regions and ribosome density define the extent of TDD

Having uncovered a global impact of mRNA translation on mRNA decay, we next investigated which mRNA features could explain the extent of TDD. To this aim, we built a random forest model to predict the observed TDDindex based on transcript-related information such as total transcript length, length of the 5’UTR, coding sequence and 3’UTR length (obtained from our PAS-Seq data), ribosome-density (obtained from our ribosome profiling data), density of m6A sites (experimentally obtained from T CD4+ mouse cells^8^) or the number of upstream Open Reading Frames (uORF), among others. The model was trained with two thirds of our dataset while the remaining third was used as a test set. In the test set, the trained model was able to capture 32% of the variance and yield a Spearman correlation coefficient of 0.58 between the predicted and observed TDDindex values (data not shown).

Analysis of the results from the random forest model led to the identification of 3’UTR length, ribosome density, and 5’UTR length as the main features predicting the observed TDDindex (Figure 4a). These results are consistent with a direct role of ribosomes in mediating decay of the mRNA they translate. To understand the relationship between these factors and the TDDindex, we plotted each of them against the observed TDDindex. Moreover, since TDD appears to be dependent on multiple transcript features, we decided to use a binning strategy to smooth the overall population and obtain a better view of the population trends (Figure 4b). Transcripts were sorted with respect to the parameter to be tested against TDDindex (for example 5’UTR length), bins of 20 adjacent transcripts were then generated and their mean TDDindex and specific transcript feature were calculated. These mean values were finally plotted against each other. Scatter-plots of 5’UTR and 3’UTR length against TDDindex indicate a clear negative correlation between these two transcripts features and the TDDindex (transcripts with either short 3’UTR or 5’UTR being more prone to TDD than those with long UTRs) (Figure 4b), thus confirming the results from the random forest analysis. Interestingly, ribosome density shows a biphasic relationship with TDDindex as we observe a strong positive correlation for low to medium ribosome density values that appears to saturate and reach a plateau for high ribosome densities (Figure 4b). Importantly, similar results are obtained when either cycloheximide (which blocks ribosomes on the coding sequence) or harringtonine (which allows 80S ribosome run-off from the coding sequence) were used to inhibit translation (Figure 4b and supplementary figure 4). This indicates that the observed relationship between ribosome density, UTR length and translation-dependent mRNA decay are not caused by a technical artifact resulting from ribosome accumulation at the coding sequence upon translational repression.

**Figure 4.**
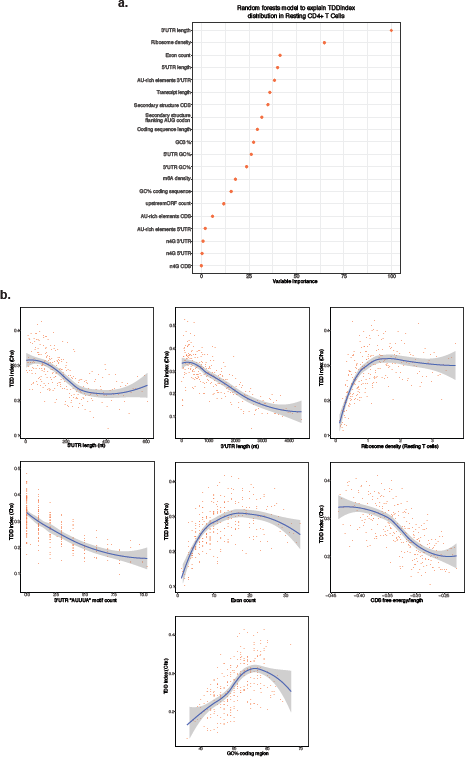
Characterization of *cis*- and *trans*-acting features linked to TDD. **a.** Random forest decision trees analysis of transcript features to explain the observed TDDindex values in resting CD4+ T lymphocytes. Features are sorted from top to bottom with respect to their importance in predicting the TDDindex. **b.** Binning plots of TDDindex against selected features. Transcripts are first ordered with respect to the feature to be compared to TDDindex and groups of 20 transcripts made along the selected feature. The mean TDDindex and feature values are plotted for each group.

Furthermore, per nucleotide predicted secondary structure in the coding region as well as occurrence of the AU-rich core pentamer motif (AUUUA) in the 3’UTR and exon count appeared as additional factors linked to the TDDindex in resting T cells (Figure 4a). AU-rich elements are able to recruit specific RNA-binding proteins involved in modulating mRNA stability^9^, however it is not clear from the literature whether AU-rich mediated decay requires on-going translation or not. Secondary structure in the coding region has been recently involved in modulating mRNA stability in a translation-dependent manner^10^, while exon count has also been linked to mRNA stability in human T lymphocytes^11^. Scatter-plots of per-nucleotide secondary structure in the CDS against the TDDindex in resting T cells (Figure 4b) indicate a global anti-correlation between the two variables (i.e. transcripts with highly structured CDS appear more prone to TDD than transcripts with less structured CDS). Occurrence of the AU-rich core pentamer motif in the 3’UTR of cellular transcripts also shows a negative correlation with the TDDindex (Figure 4b). Similarly, when looking at the extended AU-rich motif (WWWAAUUUAAWWW), which is the functional unit required to induce mRNA decay, we observe that transcripts bearing this extended motif have significantly smaller TDDindex than the overall transcript population (Supplementary figure 4b). This result suggests that AU-rich mediated mRNA decay does not occur through a translation-dependent mechanism in T cells. Finally, the number of exons per transcript displays a positive correlation with the TDDindex (Figure 4b). Yet, this link is not driven by transcript or CDS length, which both are slightly anti-correlated with the TDDindex (data not shown). This data suggests that an increased number of exon-exon junctions within a transcript appears to be linked to a higher tendency for translation-dependent mRNA decay.

Taken together, our results suggest that UTR length and density of ribosomes across the coding sequence could play a major role in mediating or modulating the extent of translation-dependent mRNA decay observed in our datasets. Nevertheless, other features such as exon count and the nucleotide composition of the coding sequence and 3’UTR could also play a significant role in mediating translation-dependent mRNA decay.

### Translation-independent mRNA decay is associated with small transcripts with few exons and a biased nucleotide composition in their coding sequence

When applied to the TIDindex, random forest analysis was able to explain 40% of the variance and yield a Spearman coefficient correlation of 0.69 between the predicted and observed TIDindex values of the test set. In this case, the number of exons per transcript as well as total transcript length were among the two most important factors able to predict the extent of TID in resting T cells (Figure 5a). In contrast to what is observed with the TDDindex, the TIDindex shows a strong anti-correlation with the number of exons per transcripts (Figure 5b). Transcripts containing few exons are strongly degraded in a translation-independent manner while transcripts with a large number of exons are poorly degraded in a translation-independent fashion. Consistent with this observation, long transcripts (which usually have a large number of exons) tend to be less dependent on TID for their decay (Figure 5b). However, among shorter transcripts, there is no clear trend and rather a small positive correlation between transcript length and the extent of TID (Figure 5b). This suggests that the effect of exon number on TID is independent from the total length of the transcript similarly to what observed for TDD.

**Figure 5.**
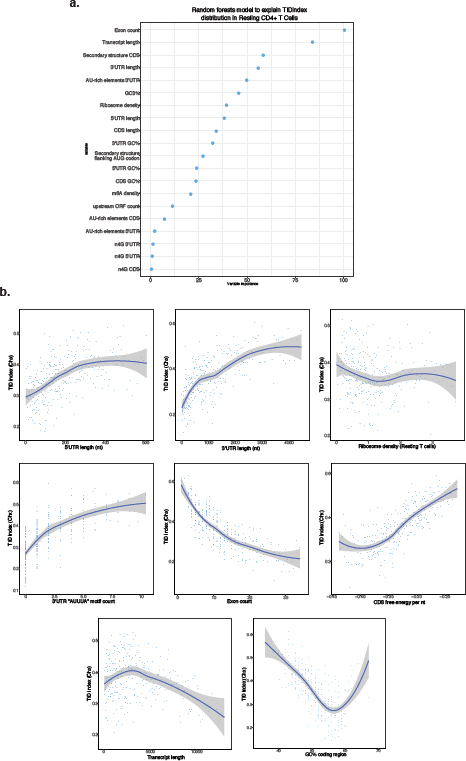
Characterization of *cis*- and *trans*-acting features linked to TID. **a.** Random forest decision trees analysis of transcript features to explain the observed TIDindex values in resting CD4+ T lymphocytes. Features are sorted from top to bottom with respect to their importance in predicting the TIDindex. **b.** Binning plots of TIDindex against selected features. Transcripts are first ordered with respect to the feature to be compared to TIDindex and groups of 20 transcripts made along the selected feature. The mean TIDindex and feature values are plotted for each group.

As observed with TDD, the length of transcript untranslated regions also appears to be linked to the extent of TID although in an opposite trend. Indeed, both in resting and activated T cells, we observe a positive correlation between 5’UTR and 3’UTR length and the extent of TID (Figure 5b). Conversely, the length of the CDS shows a similar trend to that observed with total transcript length being mildly positively correlated with the extent of TID for short transcripts and then showing a clear negative correlation for longer transcripts (data not shown). Similarly to untranslated regions, the relationship between the degree of secondary structures in the CDS and the extent of TID shows a mirror image to that observed for TDD, indicating that transcripts with highly structured coding sequences tend to be more degraded in a translation-independent manner (Figure 5b). AU-rich elements in the 3’UTR are positively correlated with the TIDindex (Figure 5b and supplementary figure 5b), further corroborating that this active decay pathway mediated by specific RNA-binding proteins is mainly translation-independent. Finally, although ribosome density is positively correlated to the extent of TDD (Figure 4b), we did not observe any particular relationship between ribosome density and the extent of TID (Figure 5b). This suggests that either ribosome density is not directly involved in modulating TID or that its effect is masked by confounding factors.

Taken together, the extent of TID appears to be controlled by similar features than TDD although in an inverted manner, further suggesting that these two pathways of mRNA decay could be extensively interconnected. TID mainly targeting transcripts with few exons and long untranslated regions.

### T cell activation reprograms the relationship between GC content in the coding region and TDD/TID

Having characterized features involved in TDD and TID in resting cells, we tested whether T cell activation could induce changes in these relationships. Indeed, upon activation, T cells undergo a dramatic change in their gene expression program (See figure 1) and metabolism to exit quiescence and enter into a proliferative state.

Random forest analysis performed with the TDDindex and TIDindex in activated cells yielded similar results to resting cells, except that secondary structure in the coding region was a less important predictor of both TDD and TID in activated cells (Supplementary figure 6a and 6b). Conversely, the GC and GC3 content in the coding region became a better predictor of both TDD and TID in activated cells (Supplementary figure 6a and 6b). Binning plots in activated cells revealed relationships between 3’UTR and 5’UTR lengths, ribosome density and the TDDindex, similar to those observed in resting cells (Supplementary figure 6c and 6d). Surprisingly, the relationship between secondary structure in the coding sequence and TDDindex or TIDindex was lost upon T cell activation (Supplementary figure 6c and 6d). Moreover, GC content in the coding region, which is positively correlated to the TDDindex and negatively correlated to the TIDindex in resting T cells (Supplementary figure 4c and 5b), displays the opposite trend upon T cell activation (being negatively correlated with the TDDindex and positively with the TIDindex, see supplementary figure 6c and 6d). Interestingly, proliferating and differentiated cells were shown to display distinct tRNA expression patterns ^12^. A change in tRNA expression has also been shown to occur during the early phases of activation of mouse T CD4+ lymphocytes^13^. Furthermore, it implicates tRNAs displaying a clear nucleotide composition bias at the first position of the anticodon loop, which corresponds to the third position of the codon they decode (“proliferation tRNAs” being enriched for codons ending with A/U, while “differentiation tRNAs” are enriched for codons ending with G/C). Finally, GC content affecting codon usage and translation efficiency has been described as an important determinant of mRNA stability^14^ thus raising the question regarding the link between TDD and GC content in the coding region.

Since our datasets allow us to discriminate between TDD and TID, we tested the relationship between codon usage and the two mRNA decay pathways. The relationship between codon usage and transcript stability has been recently studied using a specific metric known as the “Codon occurrence to mRNA Stability Correlation coefficient (CSC)”^15^. Briefly, the CSC is defined as the Pearson correlation between the frequency of each codon in mRNAs and the half-lives of the mRNAs. Here, rather than calculating a correlation with mRNA half-lives, we implemented the TDD-CSC and TID-CSC metrics by calculating the Pearson correlation between codon frequency and TDD or TID index, both in resting and activated T cells.

TDD-CSC and TID-CSC values are within similar ranges (between −0.2 and 0.2, see figure 6a and 6b) to those obtained from the literature using mRNA half-lives as input^15–17^, thus confirming the relevance of this approach to TDDindex and TIDindex. Consistent with the random forest analysis, in resting T cells, we could clearly see a nucleotide bias within codons bearing positive or negative TDD-CSC and TID-CSCs (Figure 6a and 6b, left panels). In the case of TDD, GC-rich codons are mostly associated with positive TDD-CSCs (i.e. GC-rich codons being more frequent among transcripts highly regulated by TDD), while AU-rich codons are mainly associated with negative TDD-CSCs (i.e. AU-rich codons being more frequent among transcripts poorly regulated by TDD) (Figure 6a). The opposite observation was made for TID, where GC rich codons are most frequently associated with negative TID-CSCs (i.e. GC-rich codons being more frequent among transcripts poorly regulated by TID), while AU-rich codons typically display positive TID-CSCs (i.e. AU-rich codons being more frequent among transcripts highly regulated by TID) (Figure 6b). This observation was even more pronounced when looking at the GC content of the third position of codons (GC3) (Figure 6a and 6b). Notably, for TDD, 21 out of 24 codons with a positive TDD-CSC value ended with a G or C (corresponding mainly to differentiation codons, figure 6a and 6b see blue codons), while only 10 out of 37 codons with a negative TDD-CSC value ended with a G or C (Supplementary figure 7a) leading to a significant bias in GC3 content among codons with positive and negative CSC(TDD) values (p-value=1.34e^-5^, ² test). For TID, only 6 out of 33 codons with a positive TID-CSC ended with a G or a C while only 25 out of 28 codons with a negative TID-CSC ended with G or C (Supplementary figure 7c, p-value=1.30e^-7^, X² test). Strikingly, upon T cell activation, the relationship between codon GC content, TDD and TID is flipped, both for TDD and TID, leading to GC3-rich codon being mainly associated with negative TDD-CSCs (p-value=0.0012, X² test) and positive for TID-CSCs (p-value=0.0002, X² test), while AU-rich codons (corresponding to proliferation codons, figure 6a and 6b see red codons) are mainly associated with positive TDD-CSCs and negative TID-CSCs (Figure 6a and 6b and supplementary figure 7b and 7d).

**Figure 6.**
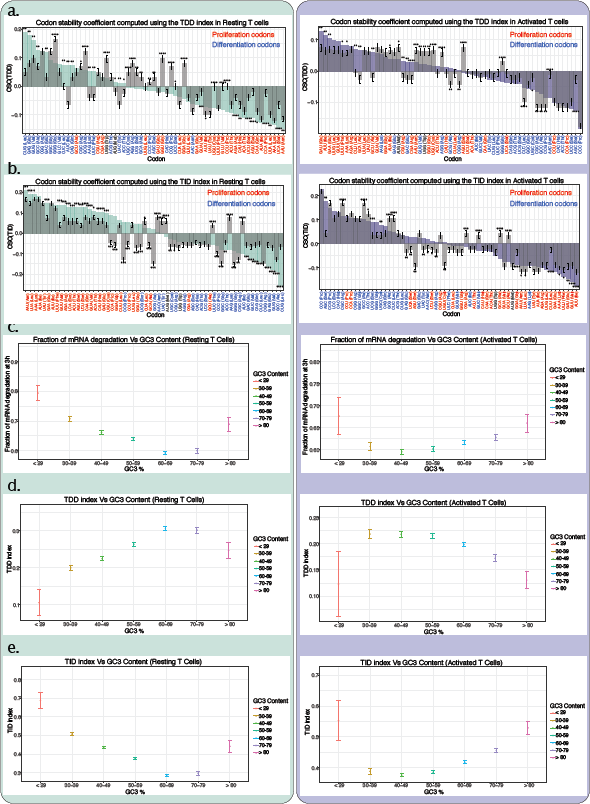
Relationship between GC content in the coding sequence, TDD and TID. **a.** Codon stability coefficient computed for each codon using the TDDindex (CSC(TDD)) in resting (left panel, green bars) and activated (right panel, violet bars) T CD4+ cells. Grey bars correspond to the mean CSC(TDD) value obtained after each coding sequence was randomized 100,000 times, maintaining the same relative proportion of each base. Observed CSC values were then compared to the distribution of randomized CSC values and a p-value calculated to test if the observed and randomized CSC values are statistically different. Codons colored in blue correspond to “differentiation codons” as described in ^13^ while codons colored in red correspond to “proliferation codons”. **b.** Same as (**a.**) but using the TIDindex (CSC(TID)) instead of the TDDindex. **c.** Plots of observed fraction of mRNA degradation (mean and standard deviation) at 3h (triptolide) against GC3 content in resting (left panel) and activated (right panel) CD4+ T lymphocytes. **d.** Same as (**c.**) but using the TDDindex instead of the fraction of mRNA degradation. **e.** Same as **c.** but using the TIDindex.

To test whether the observed CSC values are mainly driven by the nucleotide composition bias of the coding sequence, we performed permutation analysis where all coding sequences were randomly shuffled (keeping their overall nucleotide composition constant) and the CSC computed at each permutation (Figure 6a and 6b, grey bars). Then the observed CSCs were compared to the CSCs obtained from permutation and we tested whether they were significantly different. In activated T cells, the TDD-CSC and TID-CSC values obtained from nucleotide permutation followed a similar trend than those obtained from the real coding sequence (Figure 6a and 6b, right panels, compare grey bars with violet bars), arguing for an important role of GC content in driving the observed correlations, with few exceptions. Notably, all alanine codons displayed significantly higher observed TDD-CSC scores (and lower TID-CSC scores) than expected simply by the GC content of the coding region (Figure 6a and 6b, right panels). A similar observation was made for several arginine codons that displayed TDD-CSC and TID-CSC values significantly different than expected considering the GC content of the coding region. In resting T cells, however, the difference between the observed TDD-CSC, TID-CSC and the expected values from the permutation tests appears more pronounced than in activated T cells (Figure 6a and 6b, compare compare grey bars and green bars). This suggests that GC content in the coding region, alone, is not entirely responsible for the observed TDDCSC and TID-CSC values obtained for each codon. Interestingly, all proline codons (which are rich in cytidines) systematically displayed higher TDD-CSC values than those observed upon sequence permutation, possibly suggesting a role for the proline amino-acid in modulating TDD independently from the nucleotide composition of its codons (Figure 6a, left panel).

mRNA stability and GC3 content have been recently shown to be linked in human cells, where transcripts with high GC3 content were reported to display longer half-lives than those with poor GC3 content^18^. Similarly, in resting T cells, our results also indicate that transcripts with high GC3 content are more stable than those with low GC3 content (Figure 6c). As previously described^18^, we also observe a minimum of mRNA degradation at 70% GC3 content before the trend is reversed for transcripts with higher GC3 content. However, when we separate mRNA stability into translation-dependent and independent mRNA decay, we observe that these two pathways display opposite trends with respect to the GC3 content of transcripts. The TDDindex is positively correlated to GC3 content (Figure 6d) while TIDindex shows an overall negative correlation (Figure 6e), both showing a trend reversal at ~70% GC3 content similarly to what was observed with mRNA degradation. Furthermore, the relationship between TIDindex and GC3 content spans a wider range of TIDindex values than that observed for TDDindex, therefore its contribution to overall mRNA decay is more important. This explains why the relationship between the fraction of measured mRNA degradation and GC3 content follows the same trend as that of TIDindex but not that of the TDDindex.

In activated T cells, however, the relationship between GC3 content, mRNA degradation, TDDindex and TIDindex is the opposite of that observed in resting T cells (Figure 6d and 6e). Similarly to resting T cells, TID predominates over TDD in activated T cells when accounting for overall mRNA degradation and GC3 content. Among transcripts with low to moderate GC3 content (<29% and up to 49%) we observe a negative correlation between GC3 content and TIDindex (Figure 6e) or the fraction of observed mRNA degradation that reverses for transcripts above 49% GC3 (Figure 6c). Again, the opposite trend is observed for TDDindex and GC3 (Figure 6d). Nevertheless, when compared to resting T cells, the relationship of GC3 content with mRNA degradation, TID and TDD is less pronounced, spanning a narrower amplitude of TIDindex and TDDindex values.

Taken together, our results suggest that GC content in the coding sequence and particularly at the third position of codons (GC3) is an important determinant for both translation-dependent and independent mRNA decay. Moreover, the differences observed between resting and activated T cells further suggest that the relationship between GC content in the coding sequence and mRNA decay might be dependent on *trans*-acting factors such as tRNA abundance.

### TDD and TID are competing pathways defined by changes in ribosome density during T cell activation

Our results suggest that ribosomes themselves could act as triggers of translation-dependent mRNA decay (Figure 4b), the extent of which being further modulated by transcript features such as 3’UTR length, GC content in the coding region or exon number, among others. As a consequence, any change in ribosome density should lead to a corresponding change in the observed TDDindex. In order to test this, we plotted for each transcript the change in TDD index between activated and resting T cells against the corresponding fold change in ribosome density (Figure 7a). We observed a moderate but significant positive correlation (r=0.21) between changes in ribosome density and changes in TDDindex for the entire transcript population. The same observation was made whether cycloheximide or harringtonine were used to block translation in resting and activated T cells (compare Figure 7a to supplementary figure 8a). Transcripts displaying an increase in their ribosome density upon T cell activation tend to display a concomitant increase in their TDDindex while transcripts displaying a decrease in their ribosome density tend to display a decrease in their TDDindex. To test whether changes in TDDindex could also be driven by other transcript features, we applied random forest analysis to rank the features that could explain the extent of changes in both indexes. As expected, changes in ribosome density ranked among the best variables explaining changes in TDDindex, together with GC content in the coding sequence and in the 3’UTR (Supplementary Figure 8d). The fact that the GC content in the coding sequence impacts differently the TDDindex upon T cell activation agrees with our previous observation of an inversion in the relationship between TDDindex and GC3 content upon T cell activation (Figure 6d). Indeed, transcripts with high GC3 content in their CDS display a decrease in their TDDindex and transcripts with poor GC3 content in their CDS displaying an increase in their TDDindex upon T cell activation (Figure 7e). Interestingly, changes in ribosome density observed upon T cell activation are also linked to the GC3 content of their coding sequence (Figure 7f) thus suggesting that the effect of GC3 content on TDD occurs, at least partially, through the modulation of ribosome density.

**Figure 7.**
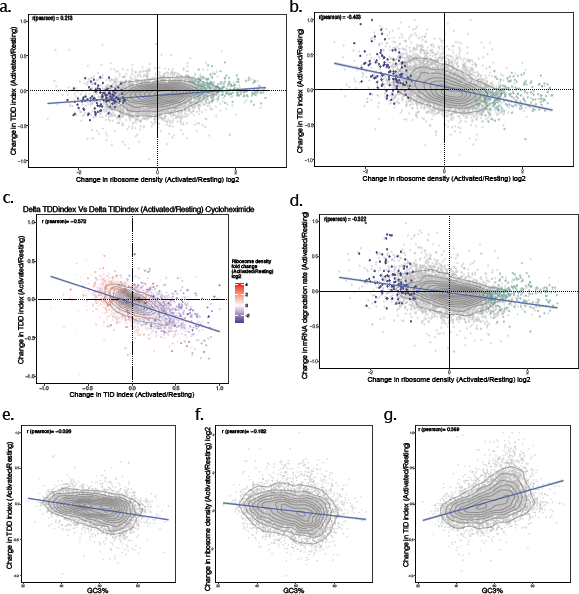
Changes in ribosome density upon T cell activation modulate both TDD and TID. **a.** Scatter plot of the changes in ribosome density between resting and activated cells (x axis) against the changes in the TDD index (y axis). Each dot corresponds to a single transcript. Violet and green dots correspond to transcripts from figure 1d displaying a significant decrease (violet) or increase (green) in ribosome density upon T cell activation. **b.** Scatter plot of the changes in ribosome density between resting and activated cells (x axis) against the changes in the TID index (y axis). **c.** Scatter plot of the change in TDDindex (y axis) and TIDindex (x axis) between resting and activated cells. Transcripts are colored with respect to the change in ribosome density measured between resting and activated cells. **d.** Scatter plot of the changes in ribosome density between resting and activated cells (x axis) against the changes in mRNA degradation rate (y axis). **e-g.** Scatter plot of the GC3% against changes in TDDindex, ribosome density and TIDindex between resting and activated cells.

Having identified changes in ribosome density as well as GC3 content in the CDS as important factors driving observed changes in TDDindex upon T cell activation, we performed a similar analysis to identify factors driving changes in the observed TIDindex. Surprisingly, changes in ribosome density were also among the main factors explaining the observed changes in TIDindex upon T cell activation. However, contrary to changes in TDDindex which are positively correlated to changes in ribosome density, changes in ribosome density display a robust negative correlation against changes in TIDindex (r=-0.403, see Figure 7b). GC content of the CDS was also the main factor explaining changes in TIDindex upon T cell activation and again, the relationship is inverted compared to that observed for TDDindex (Figure 7g). Together, these results suggest that changes in TDDindex and TIDindex upon T cell activation follow opposite trends and are, in part, defined by similar factors. Supporting this conclusion, changes in observed TDDindex between resting and activated cells are globally anti-correlated to the corresponding changes in observed TIDindex (r=-0.572) and explained by changes in ribosome density (Figure 7c). Any increase in the extent of TDD upon T cell activation generally leads to a concomitant decrease in the extent of TID and vice versa. Finally, as previously observed, the overall contribution of TID in total mRNA decay is preponderant over that of TDD. As a consequence, the fraction of mRNA degradation is negatively correlated to changes in ribosome density upon T cell activation (Figure 7d) resulting in mRNAs with increased ribosome density upon activation being stabilized while those displaying a reduction in ribosome density are less stable upon T cell activation.

Taken together, our results strongly suggest that TDD and TID occur through mutually exclusive pathways that compete for the same substrate and are defined by ribosome density and transcript *cis*-acting factors that participate in modulating the efficiency of each pathway.

## Discussion

The initial goal of this study was to investigate the hypothesis that TDD could be a key regulator of protein output during activation of immune cells. To our surprise, instead of detecting a discrete population of TDD-regulated transcripts enriched in NMD features, our results uncovered a global effect of mRNA translation in inducing mRNA decay (Figure 2c). Furthermore, the extent of TDD appeared as a transcript specific feature, with most cellular transcripts displaying moderate TDD (median of 27% of observed mRNA decay at 3h being explained by a translation-dependent mechanism), while others were either entirely degraded through TDD or completely insensitive to it. As expected, known TDD targets such as UPF2 regulated transcripts, were highly dependent on translation for their decay while lincRNAs were not (Figure 2f). By experimentally quantifying translation-dependent and independent mechanisms separately, we were further able to study each pathway independently and identify factors involved in their modulation. Overall, our results indicate that TDD and TID are governed by similar *cis*- and *trans*-acting factors, albeit in opposite manners (see Figure 8). Furthermore, upon T cell activation, changes in the extent of TDD for any given transcript is accompanied by a compensating change in the extent of TID, suggesting that these two pathways are functionally interconnected, but in a mutually exclusive manner. Altogether, our results highlight the complexity of translation-dependent and independent mRNA decay within cells and how extensively interconnected these two processes are.

**Figure 8.**
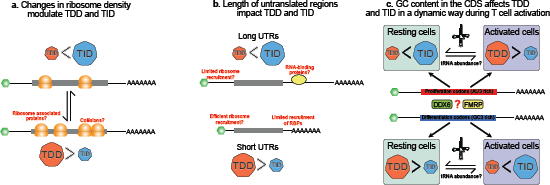
Working model presenting the main features associated with TDD and TID. **a.** Changes in ribosome density are associated with opposing changes in the extent of TDD and TID that could be explained by ribosome collisions or ribosome associated factors implicated in mediating mRNA degradation. **b.** Transcripts bearing long 5’UTR and 3’UTRs are more susceptible to TID, while transcripts with short UTRs appear more susceptible to TDD. This relationship could be explained by a higher propensity in the recruitment in the 3’UTR of RNA-binding proteins (RBPs) implicated in mRNA degradation, or a restriction of 40S ribosome recruitment for long 5’UTRs leading to TID. On the contrary mRNAs with short UTRs would recruit ribosomes more efficiently and display fewer binding sites for RBPs involved in mRNA degradation thus leading to TDD. **c.** GC content in the CDS and codon usage are important drivers of TDD and TID, possibly through specific RBPs such as DDX6 and FMRP. However, other trans-acting factors such as the abundance of tRNAs could be involved in modulating the relationship between codon usage, TDD and TID.

### Ribosome density is an important determinant for TDD

Random forest analysis of transcript features linked with TDD revealed that ribosome density is an important determinant of TDD both in resting and activated T cells (Figure 4). The relationship between TDD and ribosome density is biphasic with a positive linear correlation between TDD and ribosome density for low to medium density values, that reaches a plateau for higher ribosome densities (Figure 4b). Further confirming this relationship, changes in ribosome density upon T cell activation are positively correlated to changes in their extent of TDD (Figure 7a). Altogether, these results strongly suggest that ribosomes, themselves, are the main factor responsible for inducing TDD, most likely via recruitment of specific decay factors. This hypothesis is supported by evidence showing that ribosomes are hubs for the assembly of factors involved in mRNA surveillance and have been shown to directly interact with different mRNA decay factors^19–23^.

This scenario raises, however, several questions. Do all ribosomes have the same probability of recruiting mRNA decay factors or are there additional transcript specific determinants that condition this recruitment? Studies performed in budding yeast (*S. cerevisiae*) have pointed to codon usage coupled to ribosome residency time as features conditioning recruitment of mRNA decay factors to ribosomes and thus leading to translation-dependent decay^15,24^. Similar results have also been observed in higher eukaryotes where enrichment in specific codons were shown to lead to higher mRNA decay rates^16–18,25^. Furthermore, the frequency of ribosome collisions has been recently reported to be positively correlated to ribosome density in a transcriptome-wide manner^26^ and in some instances to recruit the RNA decay factor SKIV2L^20^, suggesting that ribosome collisions could be responsible for the correlation between ribosome-density and TDD observed in our datasets. Moreover, there is evidence showing that ribosome collisions can result in the recruitment of the ubiquitin ligase ZNF598 to the collided ribosomes, thus triggering transcript degradation^27–29^. Interestingly, ZNF598 is usually found is substoichiometric amounts compared to ribosomes^28^. This would imply that not all collision events result in mRNA degradation, thus potentially explaining the lack of a clear cleavage at stalling sites observed by Arpat and colleagues in their study^26^. Another possibility would be that ribosome collisions could induce mRNA degradation in an endonucleolytic independent manner.

### A role for untranslated regions in modulating TDD?

Our random forest analysis also pointed to 3’UTR length, and to a lesser extent to 5’UTR length, among important factors explaining TDD (Figure 4a). Further analysis confirmed a global anti-correlation between UTR length and TDD susceptibility (Figure 4b). This result is surprising since long 3’UTRs have been linked to Upf1-dependent mRNA decay^30,31^. However, there is evidence that the role of Upf1 in the decay of long 3’UTRs could occur through a ribosome-independent pathway implicating the microRNA pathway and the CCR4-NOT deadenylase complex^32^ as well as independently from NMD within chromatoid bodies through TDRD6^33^. Furthermore, in zebrafish, 3’UTR length has been shown to play a buffer role against deadenylation and decay of maternal mRNAs mediated by codon usage^34^. In this pathway, long 3’UTRs were shown to decrease accessibility of mRNA decay factors recruited by non-optimal codons to the 3’ poly(A) tail (distance model of codon-dependent mRNA decay). If a similar mechanism is responsible for protecting mRNAs from the translation-dependent mRNA decay observed in our datasets, it would imply that decay occurs through transcript deadenylation rather than an endonucleolytic cleavage.

In contrast to 3’UTR length, the role of 5’UTR length in modulating TDD is likely to be indirect, possibly through the regulation of translation initiation, which impacts ribosome density in the coding sequence. Long 5’UTRs could restrict ribosome loading and the rate of translation initiation through secondary structure or the presence of upstream ORFs and thus decrease the density of ribosomes along the CDS, leading to low TDD. Interestingly, a recent study showed that 40S ribosomes remain cap-tethered during scanning and thus only a single 40S is able to scan the 5’UTR at any given time^35^. The authors also showed that the length of the 5’UTR defines the efficiency of ribosome loading^35^. These results could explain the global anti-correlation that we observe between the length of the 5’UTR and the extent of observed TDD.

Further work will be necessary to characterize the exact mechanism by which untranslated regions could modulate TDD. Particularly for the 3’UTR which is a platform for the recruitment of RNA-binding proteins implicated in modulating translation and decay of mRNAs.

### Codon usage, GC content and tRNA availability

Studies performed in budding yeast (*S. cerevisiae*) have pointed to codon usage coupled to ribosome residency time as features conditioning recruitment of mRNA decay factors to ribosomes and thus leading to translation-dependent decay^15,24^. Similar results have also been found in higher eukaryotes were enrichment in specific codons were shown to lead to higher mRNA decay rates^16,17,25^. GC and GC3 content along the coding sequence have also been described to drive translation-dependent mRNA decay in human cells^18^ in a context where GC3-rich codons tend to destabilize mRNAs while AU3-rich codons stabilize them. Interestingly, such a GC3 bias can also be observed in the codons associated with stable and unstable mRNAs from previous work^17,25,36^, although this observation was not directly highlighted by authors.

Our strategy to discriminate between translation-dependent and independent mRNA decay allowed us to test the impact of codon usage independently on each pathway instead of relying exclusively on total mRNA degradation rates. Our results indicate that, in resting T cells, GC content in the coding sequence is indeed linked to mRNA degradation rates in a similar way as previously described, with AU3-rich transcripts being less stable than GC3-rich transcripts (Figure 6). However, when discriminating between TDD and TID, we observed that the two pathways behave in complete opposite fashion with respect to GC3 content. TDD is mainly associated with GC3-rich codons, while TID is associated with AU3-rich codons (Figure 6 a and b). Because TID, in general, accounts for a larger fraction of mRNA decay than TDD, its effects predominate when assessing mRNA degradation rates and codon usage thus introducing a bias when mRNA stability is used as a proxy to study translation-dependent mRNA decay pathways. Our results could further conciliate the impact of GC and GC3 content on mRNA stability with recent findings indicating that GC content is an important determinant for P-body localization and in defining alternative pathways for mRNA degradation^14^. According to our data, in resting cells, GC and GC3 rich transcripts are mainly degraded in a translation-dependent manner while AU and AU3 rich transcripts are mainly degraded in a translation-independent pathway. Accordingly, AU-rich transcripts are enriched within P-bodies and mainly degraded through PAT1B (although possibly outside from P-bodies^37^), while GC-rich transcripts are less abundant within P-bodies and degraded through a different pathway implicating DDX6 and Xrn1 and potentially directly linked to translation^14^. It would be therefore interesting to test the impact on TDD and TID of deleting each of these factors. Finally, as we observed a global inversion of the observed relationship between GC content in the CDS, TDD and TID upon T cell activation (Figure 6), our results further suggest that this relationship can be dynamically regulated, possibly through the modulation of trans-acting factors. Interestingly, loss of FMRP in mouse neurons leads to a reshuffling of the link between codon usage and mRNA stability^36^ similarly to the one we observe upon T cell activation. It could therefore be possible that T cell activation is accompanied by changes in the relative levels or activity of specific RNA-binding proteins involved in the recognition of specific codons or GC content in the CDS. More strikingly, it has been recently shown that tRNA abundance is dynamically regulated in mouse CD4+ T cells^13^. In this particular context, expression of tRNAs that decode “proliferation codons” bearing a strong AU3 bias tends to be up-regulated upon CD4+ T cell activation, while expression of tRNA that decode “differentiation codons” bearing a strong GC3 bias tend to be down-regulated. This observation could very well explain the switch that we observe with TDD and TID with respect to the GC3 bias in codon usage as well as the increase in ribosome density for transcripts bearing AU3-rich coding regions (Figure 7f). This result would however suggest that a rapid decoding speed would be associated with an increase in TDD.

### Ribosome loading dictates mutually exclusive and competing mRNA decay pathways?

To our surprise, changes in ribosome density upon T cell activation affected both TDD and TID but in opposite directions. Furthermore, we observe a global negative correlation (r=-0.572) in the extent of the TDD and TID changes measured upon T cell activation (Figure 7c) therefore indicating that the two decay pathways are indeed interconnected and probably competing for their mRNA substrates. In resting and activated (3h) T CD4+ lymphocytes, TID appears as more effective than TDD therefore any change towards TID tends to decrease the overall stability of the mRNA while changes towards TDD generally results in the relative stabilization of the mRNA (Figure 7d). Based on these results and the relationship between TDD/TID and GC content, which has been described as a major determinant of the targeting of mRNAs into P-bodies^14^, it is tempting to speculate that ribosome density could govern the recruitment of mRNAs to P-bodies in a dynamic manner. Supporting this hypothesis, incubation of cells with cycloheximide (which blocks elongating ribosomes on mRNAs) is known to reduce P-body size, whilst on the contrary, incubation with puromycin (which dissociates 80S ribosomes from translating mRNAs) tends to increase P-body size and abundance^38^. Conversely, changes in ribosome density could also be the consequence of an active process in which mRNAs directed towards TID transition through a translationally inactive state before being effectively degraded. MicroRNAs are an example of such a regulation as they first trigger translational repression of their target mRNA before inducing their decay. Nevertheless, although ribosome density appears to play an important role in directing mRNAs towards TDD or TID, other factors such as UTR length, exon number and GC content in the coding sequence and 3’UTR, among others, seem to play a role in further modulating the efficiency of each pathway. As a consequence, although we observe a global anti-correlation between changes in TDD and TID upon T cell activation (Figure 7c), it is nevertheless possible for both pathways to be regulated in synergy resulting in a strong stabilization or destabilization of transcripts (Figure 7c, see transcripts displaying either positive or negative changes for both TDD and TID between resting and activated cells).

### Biological roles of TDD and TID in regulating functional classes of transcripts

In resting and activated T cells, TDD principally targets transcripts coding for proteins with basic functions related to the process of gene expression, whose proteins are generally abundant in the cell. These include ribosome biogenesis factors, factors implicated in tRNA metabolism, mRNA splicing and the proteasome. For those transcripts, TDD could participate in limiting protein output per mRNA unit and introduce a negative feedback loop to avoid protein over-expression. Such a feedback loop would be particularly useful under conditions where expression is up-regulated as it is observed for several of these GO categories upon T cell activation (supplementary figure 1), when the cell is increasing in volume and requires an increase in protein levels to prepare for clonal expansion. Conversely, TID predominantly targets functional categories of transcripts that are related to the fine tuning of gene expression (transcription factors) or specific cellular pathways such as cell cycle and the immune response. These pathways require a dynamic and reversible control of gene expression and therefore mostly rely on specific trans-acting factors (such as AU-rich binding proteins) to actively regulate their expression both temporally and in their amplitude. Interestingly, transcripts highly regulated by TID display long 3’UTRs (Figure 5b) therefore providing a larger platform to recruit RBPs involved in the post-transcriptional regulation of gene expression^39,40^.

In this context, it could be of interest to identify the central players responsible for mediating TDD and TID to inactivate their function and to test the physiological consequences on T cell quiescence and activation.

## Contributions

B.C.M, M.J.M and E.P.R conceived the study and designed all experiments. B.C.M, E.P.R and A.B performed experiments, with technical assistance from L.G. E.L performed most bioinformatics analyses with help from A.C for sequencing processing of sequencing reads and codon usage calculation, D.C for the binning plot strategy and transcript database creation, F.A for G-quadruplex analysis and L.M for statistical analysis of generated data. E.P.R wrote the manuscript with contributions from all authors.

## Accession codes

High-throughput sequencing data corresponding to RNA sequencing, ribosome-profiling and poly(A)-site sequencing have been deposited in the SRA data-base under accession number GSE159301 and can be accessed upon request while the manuscript is under peer-review.

## Material and methods

### Primary cell purification and culture

Primary CD4+ T cells were obtained from 6 week old C57BL/6J female mice. Briefly, the spleen as well as the inguinal, axillary, brachial, cervical and mesenteric lymph nodes were collected, followed by ficoll separation to remove red blood cells from splenocytes. CD4+ T cells were then purified by negative selection using the CD4+ T Cell Isolation kit (Myltenyi Biotec, Cat:130-104-454) following the manufacturer’s protocol. Isolated cells were grown in RPMI medium supplemented with 10% fetal calf serum (FCS) and 50μM β-mercaptoethanol. CD4+ T cell activation was performed using magnetic beads coupled with CD3/CD28 antibodies (Thermo Fisher Scientific, Cat: 11452D) following the manufacturer’s protocol.

### Proliferation and cell surface marker detection by flow cytometry analysis

Before culture, CD4+ T cells were stained with CarboxyFluorescein Succinimidyl Ester (CFSE) (Molecular Probes). After 24, 48 or 72 h of culture, cell numbers were measured by flow cytometry using Calibrite beads (BD Pharmingen) as standards as previously described^41^. Expression of cell surface markers in resting and CD3/CD28 activated cells was made by flow cytometry using fluorescent-coupled antibodies against CD4 (Biolegend, Cat: 100434), CD3 (Biolegend, Cat: 100308), CD69 (Biolegend, Cat: 104507) and CD25 (Biolegend, Cat: 102021).

### RNA stability measurements

To monitor mRNA stability, 3 million CD4+ T cells were incubated in the presence of 5,6-Dichloro-1-β-D-ribofuranosylbenzimidazole (DRB, Sigma-Aldrich, Cat: D1916) at a final concentration of 65μM or Triptolide (Sigma-Aldrich, Cat: T3652) at a final concentration of 25μM for 15min, 1h or 3h. At each time point (0, 15min, 1h and 3h), cells were collected, counted and total RNA was extracted from the same amount of cells (3 million) at each time point using trizol in the presence of 1μl of a 1/10 dilution of ERCC Spike-In RNA (Thermo Fisher Scientific, Cat: 4456740). To monitor mRNA stability in conditions where mRNA translation is impaired, cells were incubated in the presence of either DRB or Triptolide and Cycloheximide (final concentration of 100μg.ml-1, Sigma-Aldrich, Cat: 01810) or Harringtonine (final concentration of 2μg.ml-1, Interchim, Cat: H0169) for 1 or 3h and total RNA extracted as described above. Total RNA was depleted from ribosomal RNA using Ribo-Zero rRNA Removal Kit (Human/Mouse/Rat, Illumina) followed by cDNA library preparation as described below.

### RNA-sequencing cDNA library preparation

High-throughput sequencing libraries were prepared as described in ^42^. Briefly, RNA samples depleted from ribosomal RNAs were fragmented using RNA fragmentation reagent (Ambion, Cat: AM8740) for 3 minutes and 30 seconds at 70°C followed by inactivation with the provided “Stop” buffer. Fragmented RNAs were then dephosphorylated at their 3’ end using PNK (New England Biolabs, Cat: M0201) in MES buffer (100 mM MES-NaOH, pH 5.5, 10 mM MgCl2, 10 mM β-mercaptoethanol and 300 mM NaCl) at 37 °C for 3 h.. RNA fragments with a 3′-OH were ligated to a preadenylated DNA adaptor. Following this, ligated RNAs were reverse transcribed with Superscript III (Invitrogen) with a barcoded reverse-transcription primer that anneals to the preadenylated adaptor. After reverse transcription, cDNAs were resolved in a denaturing gel (10% acrylamide and 8 M urea) for 1 h and 45 min at 35 W. Gel-purified cDNAs were then circularized with CircLigase I (Lucigen, Cat: CL4111K) and PCR-amplified with Illumina’s paired-end primers 1.0 and 2.0. PCR amplicons (12-14 cycles for RNA-seq and 4-6 cycles for ribosome profiling) were gel-purified and submitted for sequencing on the Illumina HiSeq 2500 platform.

### Ribosome profiling

Ribosome profiling libraries were prepared as described in ^43^. Cells (15 million for each biological replicate) were incubated with cycloheximide (100 μg.ml−1 final) for 5 min at 37°C. Cells were then washed two times in ice-cold PBS + cycloheximide (100 μg ml−1) and scraped in 1 ml of PBS + cycloheximide (100 μg ml−1). Cells were pelleted at 500g for 5 min at 4 °C and lysed in 1 ml of lysis buffer (10 mM Tris-HCl, pH 7.5, 5 mM MgCl2, 100 mM KCl, 1% Triton X-100, 2 mM DTT, 100 μg/ml cycloheximide and 1× Protease-Inhibitor Cocktail EDTA-free (Roche)). Lysate was homogenized with a P1000 pipette by gentle pipetting up and down for a total of eight strokes and incubated at 4 °C for 10 min. The lysate was centrifuged at 1,300g for 10 min at 4 °C, the supernatant recovered and the absorbance at 260 nm measured. For the footprinting, 5 A260 units of the cleared cell lysates were incubated with 300 units of RNase T1 (Fermentas) and 500 ng of RNase A (Ambion) for 30 min at RT. After this, samples were loaded on top of a 10–50% (w/v) linear sucrose gradient (20 mM HEPES-KOH, pH 7.4, 5 mM MgCl2, 100 mM KCl, 2 mM DTT and 100 μg ml−1 of cycloheximide) and centrifuged in a SW-40ti rotor at 35,000 r.p.m. for 2 h 40 min at 4 °C. The collected 80S fraction was complemented with SDS to 1% final and proteinase K (200 μg ml−1) and then incubated at 42 °C for 45 min. After proteinase K treatment, RNA was extracted with one volume of phenol (pH 4.5)/chloroform/isoamyl alcohol (25:24:1). The recovered aqueous phase was supplemented with 20 μg of glycogen, 300 mM sodium acetate, pH 5.2, and 10 mM MgCl2. RNA was precipitated with three volumes of 100% ethanol at −20 °C overnight. After a wash with 70% ethanol, RNA was resuspended in 5 μl of water and the 3′ ends dephosphorylated with PNK (New England BioLabs) in MES buffer (100 mM MES-NaOH, pH 5.5, 10 mM MgCl2, 10 mM β-mercaptoethanol and 300 mM NaCl) at 37 °C for 3 h. Dephosphorylated RNA footprints were then resolved on a 15% acrylamide (19:1), 8 M urea denaturing gel for 1 h 30 min at 35 W and fragments ranging from 26 nt to 32 nt size-selected from the gel. Size-selected RNAs were extracted from the gel slice by overnight nutation at RT in RNA elution buffer (300 mM NaCl, and 10 mM EDTA). The recovered aqueous phase was supplemented with 20 μg of glycogen, 300 mM sodium acetate, pH 5.2, and 10 mM MgCl2. RNA was precipitated with three volumes of 100% ethanol at −20 °C overnight. After a wash with 70% ethanol, RNA was resuspended in 5 μl of water and subjected to cDNA library construction as described above.

### Poly(A)-site sequencing

Poly(A)-site sequencing libraries were prepared as described in ^44^. Briefly, polyA?+?RNA enriched by oligoT hybridization. PolyA?+?RNA samples were then fragmented to 60–80 nt via chemical hydrolysis and reverse transcribed with anchored oligoT oligonucleotides containing forward and reverse Illumina sequencing primer sites separated by a hexaethyleneglycol spacer (Sp18) linker. At the 5′ end, each oligonucleotide began with 5′p-GG to promote ligation^42^, followed by 5 random nucleotides (unique molecular index, UMI) to enable PCR duplicate removal. Each primer also harbored a unique 5 nt Hamming barcode (BC), allowing for sample multiplexing. Following cDNA circularization with CircLigase I, libraries were PCR amplified (12–14 cycles) and subjected to single end 100 nt sequencing on Illumina’s Hiseq platform.

### Analysis of high-throughput sequencing reads

All the scripts used for this analysis are available at the following repository https://gitbio.ens-lyon.fr/LBMC/RMI2/

Sequencing reads were split with respect to their 5′ in-line barcode sequence. After this, 5′-barcode and 3′-adaptor sequences were removed from reads using FASTX-Toolkit (http://hannonlab.cshl.edu/fastx_toolkit/index.html). Reads were mapped to a custom set of sequences including mouse 18S, 28S, 45S, 5S, and 5.8S rRNA, tRNAs and the ERCC Spike-In sequences using Bowtie/2.2.4 ^45^ with the following parameters “bowtie2 -t --fast”.

For RNA sequencing samples, reads that failed to map to this custom set of sequences were next aligned to the mouse mm10 assembly and the gencode vM7 annotation using TopHat 2 (v2.0.13) ^46^ with the following parameters “tophat2 --bowtie1 (bowtie version 1.1.1.0) --library-type fr-secondstrand --b2-sensitive -i 30 -m 1 -g 10 --max-coverage-intron 1000000”. Read counts on all transcripts of interest were obtained using the HTSeq count package ^47^ with the following parameters “htseq-count -f sam -r pos -s yes -a 10 -m intersection-nonempty”.

For ribosome profiling samples, reads that failed to map to this custom set of sequences were next aligned to the mouse mm10 assembly and the gencode vM7 annotation using TopHat 2 (v2.0.13) ^46^ with the following parameters “tophat2 --library-type fr-secondstrand --b2-sensitive -i 30 -m 1 -g 10 --max-coverage-intron 1000000”. Read counts on gencode vM7 appris principal transcripts ^1^ were obtained using the HTSeq count package ^47^ with the following parameters “htseq-count -f sam -r pos -s yes -a 10 -m intersection-nonempty”.

### Transcript database creation

The mouse transcript database was generated using the genome assembly and sequences files gencode.vM23.annotation.gff3 and GRCm38.p6.genome.fa obtained from https://www.gencodegenes.org/. A SQLite database was first generated using the python package GffUtils (https://github.com/daler/gffutils). Then a csv file was generated for each transcript with an associated coding sequence in order to collect all available components or attributes (UTRs, CDS, introns, exons, associated gene information) and their basic properties (genomic start/end points, sequence, length, GC percentage). When available, the APPRIS level of each transcript was also recovered.

To avoid retaining transcripts with aberrant features, an initial quality control was performed. The quality control primary filters included the presence of a canonical initiation (ATG, or CTG) and termination (TAA, TAG, or TGA) codons in the 5’ and 3’ extremities of the CDS sequence respectively. Moreover the absence of mis-sequenced regions and the length of the CDS (length > 200) were monitored. A last filter validated that the splicing donor and acceptor sites have the proper canonical extremities -GT and AG – respectively. All available experimental data (ribo-density, m6A-seq, PAS-seq) or bio-informatics analysis (TDD indexes, G-quadruplexes, specific codon stretches), were then attributed to the corresponding transcripts files. When data was only associated to the gene, all corresponding transcripts files were updated. The PASeq peaks genomic positions and counts were identified in triplicates for both cell status: Resting and Activated and already associated to a specific gene. Thus the peaks and their relative counts in each experimental condition were associated to the most suitable transcripts of the corresponding gene (for example: in an exon *vs* an intron, or to the closest 3’UTR end when downstream). Depending on the number of PASeq peaks attributed to a transcript and their positions and counts a weighted experimental 3’UTR length was then possible to calculate for each experimental condition.

As the RNAseq approach attributes reads to genes, the best transcript for each experimental condition was determined. In this purpose all the transcripts of a single gene were ranked on 4 criteria: pass the initial quality control, APPRIS level, number of PASeq peaks (*i.e.* present in the experimental condition), and length.

### Differential analysis with DESeq2

Differential expressed genes and translated genes upon cell activation were obtained using DESeq2^4^ (version 1.24.0) in R (version 3.6.3) using respectively RNAseq and Ribosome Profiling read counts of activated T cells versus resting Tcells with a classical model design ~ condition + replicate.

To obtain genes that differ in ribosome density, a DESeq2 model was constructed using both RNAseq and Ribosome profiling read counts with the following model: ~ condition:sequencing + condition + replicate:sequencing, where condition refers to the resting/activated state of the cells and sequencing to RNAseq/RiboSeq libraries. Specifically changes in ribosome density were recovered using the contrast list (“conditionactivated.sequencingRiboSeq”, “conditionresting.sequencingRiboSeq”) of the results command.

### Read counts normalization by DESeq with endogenous stable genes

To define endogenous stable genes, read counts of 0h Triptolide versus 3h Triptolide in RPM (read per million) were plotted. As the relative amount of stable gene reads increased upon blocking transcription when compared to the basal condition, stable genes were those that responded to the following condition in all three replicates (Supl. Fig 2.B): RPM3h Triptolide – 0.3 x RPM3h Triptolide < RPM0h Triptolide < RPM3h Triptolide + 0.9 x RPM3h Triptolide

Stable genes were then used by DESeq2^4^ (1.24.0) to estimate size factor of all libraries and then normalized read counts were recovered in a matrix for downstream analysis and calculation.

### Functional analysis upon cell activation

Functional enrichment analysis was performed with David^48,49^ (version 6.8, https://david.ncifcrf.gov/summary.jsp) selecting genes with an adjusted p-value < 0.05 in the differential gene expression and differential translation analysis. Plots were generated with Revigo^50^ (/http://revigo.irb.hr/) using as inputs GO Terms with an adjusted p-value < 0.01 as calculated by the David software.

### Random Forest models

First, the gene-set of interest was reduced to the expressed genes based on the gene normalized read counts of 3h Triptolide libraries (~5000 genes in resting and ~6000 genes in activated T-cells). Of these, only genes with completed observations in all tested parameters were kept. Dataset was then divided in a train and validation dataset, composed of respectively 70 % and 30 % genes randomly chose. Regression Random Forest models were built using R (version) with the caret package (version) using the command `train(predict ~., data = train_set, method = “ranger”, importance = ‘impurity’)`, where `“predict”` refers to the predicted parameters (TDDindex or TIDindex). Briefly, the train function re-runs the model over 25 bootstrap samples and across 3 options of the number of randomly selected predictors at each cut in the tree (mtry parameter).

The accuracy of the final model obtained by the train function was then verified by predicting the parameters of interest (TDDindex or TIDindex) of the validation dataset with the random forest model and calculating the Spearman correlation between the predicted versus the real value.

### G-quadruplex prediction

G-quadruplex structures within transcripts were predicted as described in ^51^.

### CSC correlation – permutation sequence

CSC scores were calculated inspired by ^15^. Briefly, the CSC score is the Pearson correlation between the frequency of occurrence of each codon and the TDDindex of the corresponding transcript. Statistical significance was determined permutating 10,000 times the sequence of each coding sequence. This allow us to generate 10,000 transcriptomes with random codon but with transcripts that share same features as the original sequence (GC content, CDS length) For each random transcriptome, the CSC score for each codon was computed allowing us to calculate a CSC score distribution obtained randomly. So, the p-value correspond to the frequency of the real CSC score obtained with the random transcriptomes. False Discovery Rates (FDR) were calculated with the Benjamini & Yekutieli method^52^ to adjust the p-value for multiple comparison tests.

### Gene ontology analysis using TDDindex and TIDindex values

Transcripts were associated to Gene Ontology terms using the mgi.gaf file (http://current.geneontology.org/annotations/mgi.gaf.gz), the GO terms to GO phrase association as well as the GO tree were obtained from the go-basic.obo file (http://purl.obolibrary.org/obo/go/go-basic.obo). The distributions of TDDindex and TIDindex values was generated for each GO and compared to the distribution obtained for the global transcript population. The statistical significance of the difference in the mean TDDindex or TIDindex value between the transcripts from any given GO and the global population was determined by a bootstrapping test (50000 random ensembles of same dimension of the GOs). The obtained p-values were then submitted to a hierarchical false discovery rate–controlling methodology [Yekutieli], using the structure of the GO tree.

## Acknowledgments

We thank Wendy Gilbert for helpful comments during the 2019 RNA meeting. We also thank Vincent Vanoosthuyse, Mélanie Wencker, Christelle Morris and RMI2 lab members for manuscript proofreading. We gratefully acknowledge support from the PSMN (Pôle Scientifique de Modélisation Numérique) of the ENS de Lyon for the computing resources. We thank the members of the LBMC biocomputing center for their involvement in this project.

This work was funded by the European Research Council (ERC-StG-LS6-805500 to E.P.R) under the European Union’s Horizon 2020 research and innovation programs, by Fondation FINOVI (to E.P.R) and by the Howard Hughes Medical Institute (to M.J.M).

---

**Supplementary figure 1.**
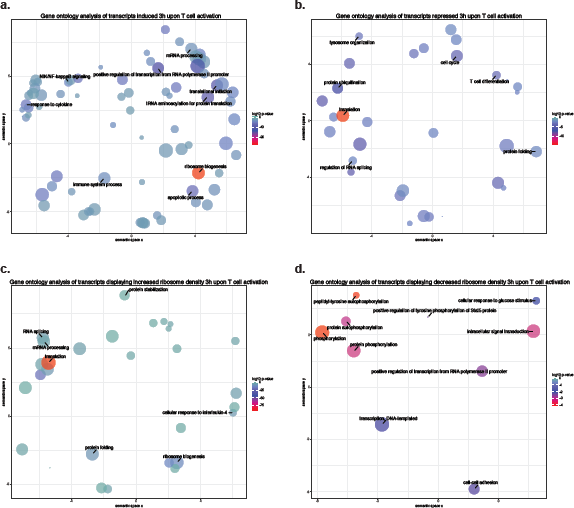
Gene-ontology analysis of differentially expressed and translated transcripts upon CD4+ T cell activation. Analysis of enriched functional gene categories among **a.** Transcripts whose expression is up-regulated upon T cell activation; **b.** Transcripts whose expression is down-regulated upon T cell activation; **c.** Transcripts whose ribosome density increases upon T cell activation; **d.** Transcripts whose ribosome density decreases upon T cell activation. Analysis were performed using REVIGO^50^. Each circle corresponds to a given gene-ontology category, the radius being linked to its size (in gene number) and the color corresponding to the adjusted p-value of the observed enrichment. The *x* and *y* axes correspond to arbitrary positions defined by REVIGO to facilitate separation and reading of enriched GO terms.

**Supplementary figure 2.**
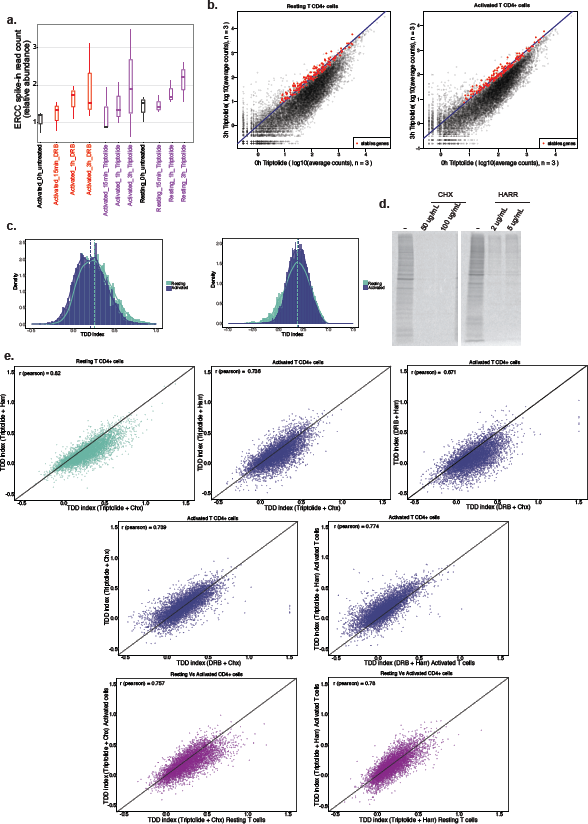
Calculation and comparison of TDD and TID indexes using different combinations of transcription and translation inhibitors in CD4+ T cells. **a.** Relative abundance of ERCC exogenous Spike-In in RNA-seq libraries prepared from cells incubated for different times with transcription inhibitors. **b.** Scatter-plots of read counts in control cells and cells incubated for 3 hours in the presence of triptolide. Red dots correspond to the stable transcripts that were selected to normalize gene expression upon transcription inhibition. **c.** Distribution of TDDindex (left panel) and TID index (right panel) in resting and activated CD4+ T cells. **d.** Metabolic labeling of resting CD4+ T cells using S^35^ methionine in the absence and presence of different doses of cycloheximide and harringtonine. **e.** Scatter-plots of TDDindex obtained with different combinations of transcription and translation inhibitors and in resting or activated CD4+ T cells. **f.** Scatter plot of TDDindex against TID index in resting (left panel) and activated (right panel) CD4+ T cells.

**Supplementary figure 3.**
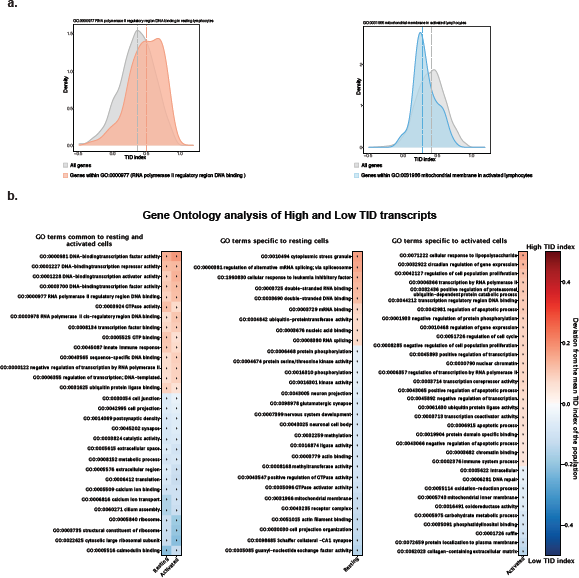
Gene ontology analysis of transcripts displaying high or low TIDindex values. **a. (Left panel)** TIDindex distribution for all expressed genes (grey) or genes from the “RNA polymerase II regulatory region DNA binding” GO category (red). **(Right panel)** TIDindex distribution for all expressed genes (grey) or genes from the “Mitochondrial membrane” GO category (blue). **b.** List of GO terms with a median TIDindex that is significantly higher (red) or lower (blue) from the median TIDindex of the entire transcript population that are common to resting and activated T cells (left panel) or specific to resting (middle panel) or activated (right panel) T cells.

**Supplementary figure 4.**
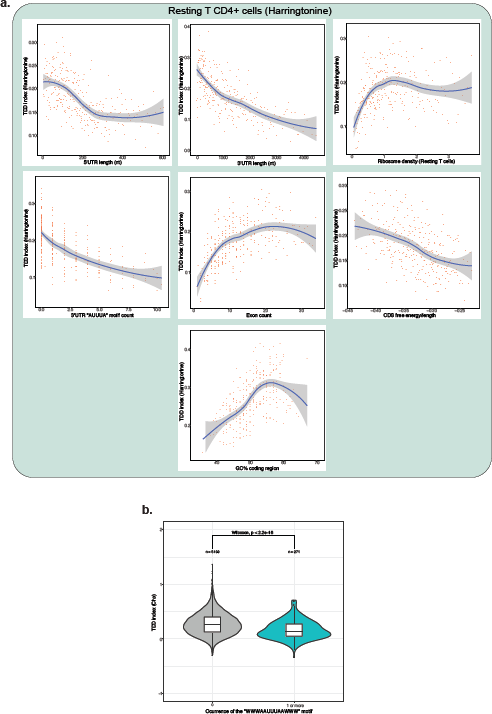
**a.** Binning plots of TDDindex obtained using harringtonine, instead of cycloheximide, against different transcript features. **b.** Violin-plot of TDDindex obtained with cycloheximide for transcripts with no “WWWAAUUUAAWWW” motif or 1 or more motifs in their 3’UTR.

**Supplementary figure 5.**
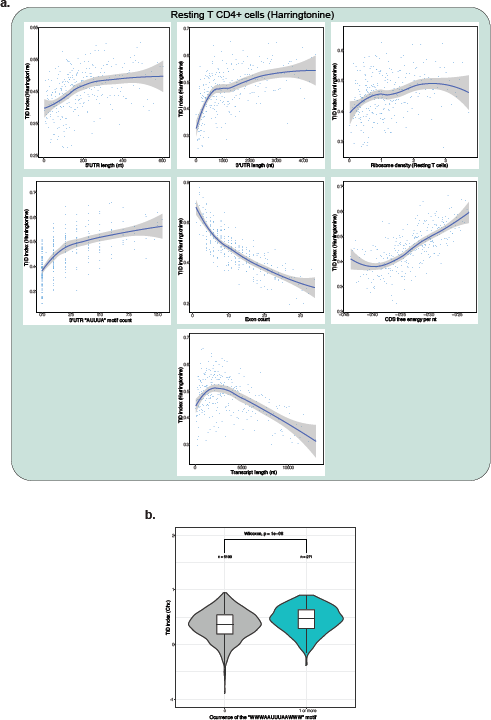
Binning plots of TIDindex obtained using harringtonine, instead of cycloheximide, against different transcript features. **b.** Violin-plot of TIDindex obtained with cycloheximide for transcripts with no “WWWAAUUUAAWWW” motif or 1 or more motifs in their 3’UTR.

**Supplementary figure 6.**
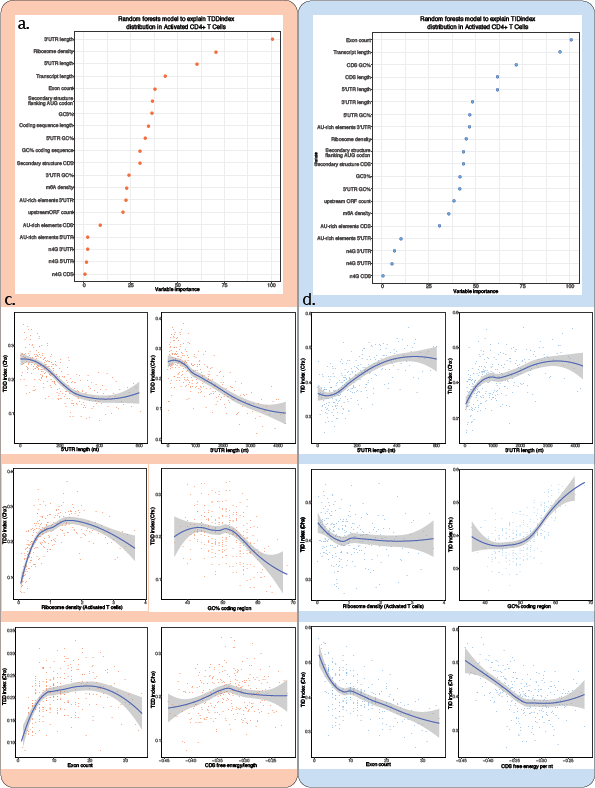
**a and b.** Random forest decision trees analysis of transcript features to explain the observed TDDindex (left panel) and TIDindex (right panel) values in activated CD4+ T lymphocytes. Features are sorted from top to bottom with respect to their importance in predicting the TDDindex or TIDindex. **c and d.** Binning plots of TDDindex (left panels) or TIDindex (right panels) against selected features. Transcripts are first ordered with respect to the feature to be compared to TIDindex and groups of 20 transcripts made along the selected feature. The mean TIDindex and feature values are plotted for each group.

**Supplementary figure 7.**
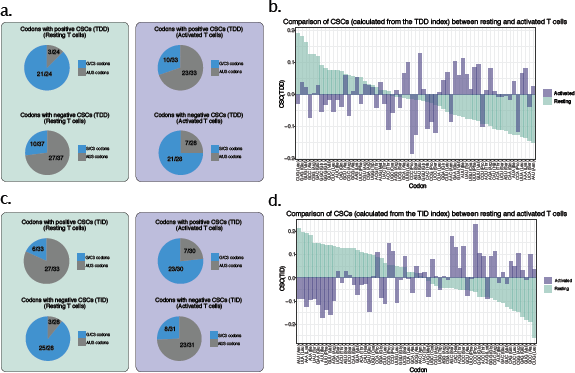
Relationship between GC content in the coding sequence, TDD and TID. **a.** Relative abundance of GC3 and AU3 codons among codons with positive and negative CSCs in resting (left panel) and activated (right panel) CD4+ T cells. **b.** Comparison of observed codon stability coefficient for the TDDindex in resting (green bars) and activated (violet bars) T CD4+ lymphocytes. **c.** Same as (**a.**) but using TIDindex instead of TDDindex. **d.** Same as (**b.**) but using TIDindex instead of TDDindex.

**Supplementary figure 8.**
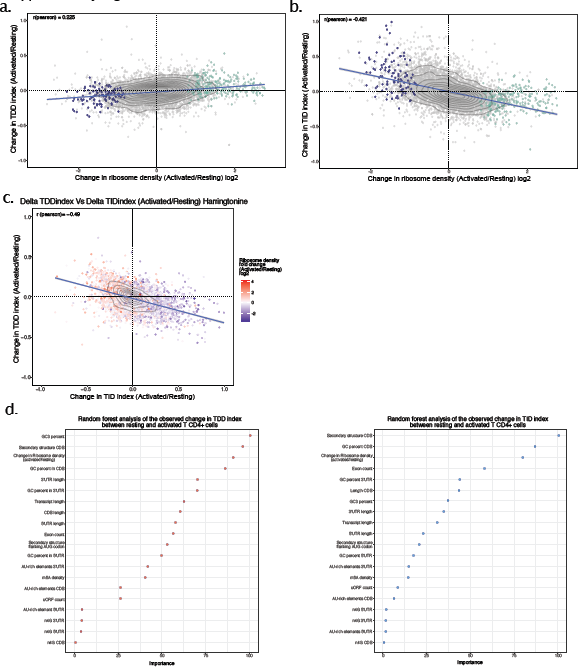
Changes in ribosome density upon T cell activation modulate both TDD and TID. **a.** Scatter plot of the changes in ribosome density between resting and activated cells (x axis) against the changes in the TDD index (y axis) obtained using harringtonine instead of cycloheximide. Each dot corresponds to a single transcript **b.** Scatter plot of the changes in ribosome density between resting and activated cells (x axis) against the changes in the TID index (y axis) obtained using harringtonine instead of cycloheximide. **c.** Scatter plot of the change in TDDindex (y axis) and TIDindex (x axis) between resting and activated cells obtained using harringtonine. Transcripts are colored with respect to the change in ribosome density measured between resting and activated cells. **d.** Random forest analysis of the changes in TDDindex (left panel) and TIDindex (right panel) between resting and activated CD4+ T cells.

**Supplementary table 1. Differential gene expression and translation analysis between resting and activated CD4 T cells; and gene ontology analysis of differentially expressed and translated genes between resting and activated CD4 T cells**.

